# Ocular pigmentation in humans, great apes, and gibbons is not suggestive of communicative functions - having an eye on the ‘cooperative eye hypothesis’

**DOI:** 10.1101/2021.03.18.435993

**Authors:** Kai R. Caspar, Marco Biggemann, Thomas Geissmann, Sabine Begall

## Abstract

Pigmentation patterns of the visible part of the eyeball, encompassing the iris and portions of the sclera, have been discussed to be linked to social cognition in primates. In the context of the *cooperative eye hypothesis*, the white sclera of humans has been viewed as a derived adaptive trait, enhancing communication via glance cueing. Here, we provide a comparative analysis of ocular pigmentation patterns in 15 species of hominoids (humans, great apes & gibbons) representing all extant ape genera, based on photographs and literature data. Additionally, we quantify hominoid scleral exposure on the genus level during different glancing situations. Our data reveals a continuum of eye pigmentation traits among the studied taxa. Gibbons display darker, more uniformly colored eyes than great apes and expose less sclera, particularly during averted glancing. Iridoscleral contrasts in orangutans and gorillas approach the human condition but differ between congeneric species. Contrary to recent discussions, we found chimpanzee eyes to exhibit a cryptic coloration scheme that resembles gibbons more than other great apes and that does not enhance glance cueing or gaze following. We critically evaluate the evidence for links between social cognition and eye pigmentation in primates, concluding that the *cooperative eye hypothesis* cannot convincingly explain the patterns observed. Although the human eye exhibits unique traits that are likely linked to social communication, high iridoscleral contrast is not one of them. Differences in scleral pigmentation between great apes and humans are gradual and might have arisen via genetic drift and sexual selection.

## Introduction

Eyes are importantly involved in human non-verbal communication (Langton et al., 2000). Glancing (i.e., eye orientation/eye gaze; opposed to gazing, i.e., head orientation) in particular can facilitate social communication, for instance as a referential cue, informing observers about one’s attentional focus (Povinelli & Eddy, 1996; Tomasello et al., 2007). It is commonly assumed that eye-mediated communication is far more sophisticated in humans than it is in other primates, if present there at all. In an influential paper, Kobayashi and Kohshima (2001) argued that this difference in communicative behavior is mirrored by the morphology of the human eye, a hypothesis popularized beforehand by Morris (1985). Humans exhibit an almost complete depigmentation of the sclera and overlying conjunctiva, creating the white of the eye, which contrasts with the darker iris. Beside its conspicuous coloration, large portions of scleral surface are exposed in the human eye due to its marked horizontally extended outline. This characteristic might have originally evolved to facilitate wide-angle glancing and thereby to extend humans’ visual field in terrestrial habitats (Kobayashi & Koshima, 2001). Kobayashi and Kohshima (2001) emphasized how ocular morphology differs between humans and non-human apes (from here on “apes”) and argued that the traits of the human eye would be uniquely suited to enable effective glance-based communication. This idea was further developed and coined the *cooperative eye hypothesis* by Tomasello et al. (2007) who characterized the human eye as a social tool to convey intentions, guide actions and mediate joint attention (see also Hare, 2017). Darker, less conspicuous eyes on the other hand would conceal glance direction in order to mask intentions, which was hypothesized to be advantageous in the more competitive social environments assumed for apes (Kobayashi & Kohshima, 2001; Perea García et al., 2019). Given that, human and non-human primate pigmentation patterns would serve contrary adaptive purposes. However, the assumption of such a clear dichotomy between ape and human eyes has been contested.

It has become increasingly apparent that ocular pigmentation in most great ape species is far more variable than assumed by Kobayashi and Kohshima (2001), who predominately studied few individuals per species in their sample (e. g. n = 2 for bonobos, n = 5 for orangutans). In Western gorillas (*Gorilla gorilla*) and bonobos (*Pan paniscus*), scleral pigmentation appears to be particularly plastic, ranging from plain black to fully white (Mayhew & Gómez, 2015; Perea García et al., 2019). A small-scale study also found predominantly light sclerae in Sumatran orangutans (*Pongo abelii*) (Perea García, 2016). Bornean orangutans (*Pongo pygmaeus*), chimpanzees (*Pan troglodytes*) and Eastern gorillas (*Gorilla beringei*) were instead reported to almost consistently display dark sclerae (Mayhew & Gómez, 2015; Perea García, 2016; Perea García et al., 2019; but see e.g., Goodall, 1986) for exceptions), so that great ape scleral coloration does not appear to follow a clear phylogenetic pattern. Additionally, it has been shown that the amount of visible sclera does not differ significantly between gorillas and humans in averted gaze situations which are of particular communicative value, demonstrating a greater than previously assumed continuity in this ocular trait as well (Mayhew & Gómez, 2015). In orangutans, scleral exposure during averted glancing is markedly lower, approximating that of humans during direct glancing (Kaplan & Rogers, 2002). Data on scleral exposure in varying gaze situations is so far lacking for other primates.

In a recent study, Perea García et al. (2019) revisited the topic of ape and human eye coloration by comparatively quantifying iridoscleral contrasts in humans, bonobos and chimpanzees. Despite pronounced differences in ocular coloration, the grayscale brightness of the iris when compared to the sclera (relative iris luminance = RIL) did not differ significantly between the studied species. Chimpanzees have light, amber-colored irises which contrast with their typically black sclerae, inversing the pattern found in humans and the majority of bonobos. Perea García et al. (2019) state that owing to comparable RIL values, ocular pigmentation patterns in the three species are equally conspicuous and suggest that chimpanzees, bonobos and humans share “cooperative eyes”.

However, there is only very little experimental support for the hypothesis that great apes or primates in general rely on conspecifics’ glancing as a communicative cue (compare Bethell, Vick, and Bard (2007)). Laboratory studies on rhesus macaques (*Macaca mulatta*) monitoring eye motion showed that these monkeys respond to conspecific glancing by reflexively aligning their own field of vision, just like humans do (Deaner & Platt, 2003). By now, comparable experimental data on apes are not available but it has been shown that chimpanzees can respond to human glances in a similar fashion (Povinelli & Eddy, 1996). Therefore, reflexive glance following can be expected to be shared by a variety of primates, although it remains speculative to which degree this information is used by them in social situations. For example, chimpanzees typically fail to exploit glancing as a referential cue in forced-choice tasks despite their ability to reflexively follow human glances (Povinelli & Eddy, 1996; Barth et al., 2005; but see Tomasello et al., 2007). In any case, it is obvious how different patterns of ocular contrast and scleral exposure might facilitate providing glance cues in a variety of situations if a respective species is able to decode them as referential cues (Perea García, 2016).

Apart from that, Perea García et al. (2019) hypothesize the depigmented sclera of humans and bonobos to be an evolutionary byproduct for selection against aggression. In that, it would mirror the pleiotropy-induced domestication syndrome described for companion animals (Cieslak et al., 2011; see also Lord et al., 2020 and Wright et al., 2020). They also imply that such selection against aggression might explain the divergent scleral pigmentation patterns among gorilla and orangutan species, suggesting a link between social patterns and ocular pigmentation. Although this byproduct hypothesis contradicts the assumption that communicative demands chiefly drive the evolution of eye color in apes, it is compatible with depigmented eyes being particularly effective in communication. Still, limited data make it hard to convincingly state a correlation between ocular pigmentation, particularly RIL, and sociocognitive functions.

To clarify this matter, it would be desirable to place ocular pigmentation in great apes into a broader evolutionary perspective. Of particular interest for such comparisons are the small apes or gibbons (family Hylobatidae, genera *Hoolock, Hylobates, Nomascus, Symphalangus*), which form the sister group to the human and great ape clade (family Hominidae). Different from great apes, gibbons are morphologically, ecologically, and socially rather uniform. All species are specialized canopy-dwellers living in small family groups (Reichard et al., 2016). Despite their phylogenetic position and resulting relevance to understand the evolution of hominoid cognitive traits, gibbons are widely ignored in primate behavioral research (excluding acoustic communication), so that little is known about their sociocognitive traits (Butler & Suddendorf, 2014; ManyPrimates et al., 2019). However, there currently is consensus that glancing does not carry noteworthy communicative value to them (Caspar et al., 2018; Sanchez-Amaro et al., 2020). In line with this, gibbon gaze following is less sophisticated than in their large-bodied relatives. Small apes follow the head orientation of both conspecifics and humans but do so in a reflexive way, while great apes deduce referential information from gaze, as indicated by double checking and habituation to repeated gazing events (Liebal & Kaminski, 2012). A comparison of great and small apes could therefore aid in identifying correlates of eye-mediated communication.

As described above, it has been argued that eye coloration in the genera *Homo* and *Pan* represents an adaptation for optimized glance cueing (Perea García et al., 2019) and that ocular coloration is tightly correlated with sociocognitive functions among great apes and humans (Kobayashi & Kohshima, 2001; Tomasello et al., 2007; Perea García, 2016). This notion implies that primates that are not expected to utilize glance cues should show less conspicuous ocular contrasts, representing an ancestral state compared to the optimized patterns (Perea García, 2016). Gibbon eyes indeed appear unspecialized based on a small mixed-species dataset presented by Kobayashi and Kohshima (2001). Compared to great apes, scleral portions of the gibbon eyeball remain largely unexposed during direct glancing, as their eyes show a circular rather than an elliptical lid excision (Kobayashi & Kohshima, 2001). The uniformly colored iris is filling out the visible part of the eye almost completely. Potential iridoscleral contrast, a trait not previously quantified in gibbons, is therefore significantly concealed. Instead, conspicuous facial fur patterns are present in many gibbon species that might effectively indicate head but not eye orientation (Geissmann, 2003).

Here, we provide data on eye contours as well as ocular pigmentation for a large sample of hylobatids to draw quantitative comparisons with hominids. For this, we also quantified iridoscleral contrasts in gorillas and orangutans, providing the first comprehensive overview of ocular pigmentation patterns across the hominoid radiation. Following the *cooperative eye hypothesis*, we predicted that gibbon eyes would be less conspicuous than hominid eyes. We further expected to find uniform patterns of ocular coloration among small ape species. Finally, we assessed eye contours and the amounts of sclera exposed during forward and averted glancing in three gibbon genera and each hominid genus. Again, we hypothesized to find consistent differences between small and great apes based on the preliminary results of Kobayashi and Kohshima (2001), with the small apes exposing less sclera than their large-bodied relatives.

## Materials and Methods

We conducted an extensive internet search for pictures of great apes and gibbons from wild as well as captive environments and pooled images found online with private digital photographs (see Suppl. Tab. 1). In case of data gathered online, information on the identity, sex and location of photographed apes were derived from the source websites to avoid repeated sampling of particular individuals. If available, international studbooks were used to check whether zoo-housed individuals were born in the wild or in captivity. Species level classification and nomenclature were adopted from Burgin et al. (2020). Within hylobatids, we pooled data from recently diverging populations to reach larger sample sizes for the respective groups. As a consequence, *Nomascus siki* was grouped under *N. leucogenys, N. annamensis* under *N. gabriellae, Hoolock tianxing* under *Ho. leuconedys*, and *Hylobates abbotti* and *H. funereus* under *H. muelleri* (see Suppl. Tabs. 1 and 2 for taxonomic identities of each subject). Since ocular pigmentation and the shape of the eye contour can differ between juvenile and mature apes, we restricted our sampling to pictures of adults. Subadult animals were only included for some gibbon species but only in case they already developed adult pelage traits (e. g., after entering the pale color phase in female gibbons of the genera *Hoolock* and *Nomascus*).

We followed the methodology of Mayhew and Gómez (2015), Perea García (2016), and Perea García et al. (2019) in using ImageJ (Schneider et al., 2012) to take quantitative measurements from the digital images we gathered.

### Quantifying ocular luminance in great and small apes

Ocular pigmentation approximated by luminance was quantified for gorillas (*G. beringei*, n = 22; *G. gorilla*, n = 40), orangutans (*P. abelii*, n = 23; *P. pygmaeus*, n = 24), crested gibbons (*N. gabriellae*, n = 20; *N. leucogenys*, n = 27), dwarf gibbons (*Hy. lar*, n = 30; *Hy. moloch*, n = 18; *Hy. muelleri*, n = 17; *Hy. pileatus*, n = 17), hoolock gibbons (*Ho. leuconedys*, n = 10), and siamangs (*S. syndactylus*, n = 29), summing up to 277 individuals (Tab. 1). Additional data on ocular pigmentation traits in adult bonobos, chimpanzees and humans was derived from Perea García et al. (2019), resulting in a total of seven hominid and eight hylobatid species representing all of the eight genera of extant apes (n_total_ = 386).

**Table 1:** Summary of data on ape ocular pigmentation patterns. Type I phenotype describes eyes in which the sclera is lighter than the iris. HC and RIL correspond to species means. * Data derive from Perea García et al. (2019); ** n = 47.

| Species | n | % of type I phenotype | HC | RIL |
| --- | --- | --- | --- | --- |
| *Hoolock leuconedys* | 10 | 0 | 30.5 | 41.8 |
| *Hylobates lar* | 30 | 46 | 34.5 | 51.2 |
| *Hylobates moloch* | 18 | 0 | 39.0 | 33.5 |
| *Hylobates muelleri* | 17 | 0 | 41.5 | 38.0 |
| *Hylobates pileatus* | 17 | 12 | 30.2 | 53.4 |
| *Nomascus gabriellae* | 20 | 30 | 17.3 | 62.0 |
| *Nomascus leucogenys* | 27 | 0 | 24.1 | 52.7 |
| *Symphalangus syndactylus* | 29 | 38 | 31.2 | 59.4 |
| *Gorilla beringei* | 22 | 41 | 56.7 | 40.2 |
| *Gorilla gorilla* | 40 | 85 | 81.7 | 35.2 |
| *Homo sapiens* * | 49 | 100 | 84.4** | 48.7 |
| *Pan paniscus* * | 24 | 88 | 61.4 | 46.4 |
| *Pan troglodytes* * | 36 | 11 | 33.3 | 48.0 |
| *Pongo abelii* | 23 | 87 | 83.7 | 38.6 |
| *Pongo pygmaeus* | 24 | 79 | 48.0 | 45.6 |

To be included into the sample, picture resolution had to be high enough to unequivocally distinguish between pupil, iris, and sclera in at least one eye of the photographed subject. We noticed that all gibbon species as well as orangutans and gorillas typically display a thin grey line encircling the peripheral iris, which appears as a salient demarcation to the sclera (see Fig. 1). We only sampled pictures on which this demarcation line was visible. In case the aforementioned criteria were met, pictures capturing both direct and averted gaze were included.

**Figure 1:**
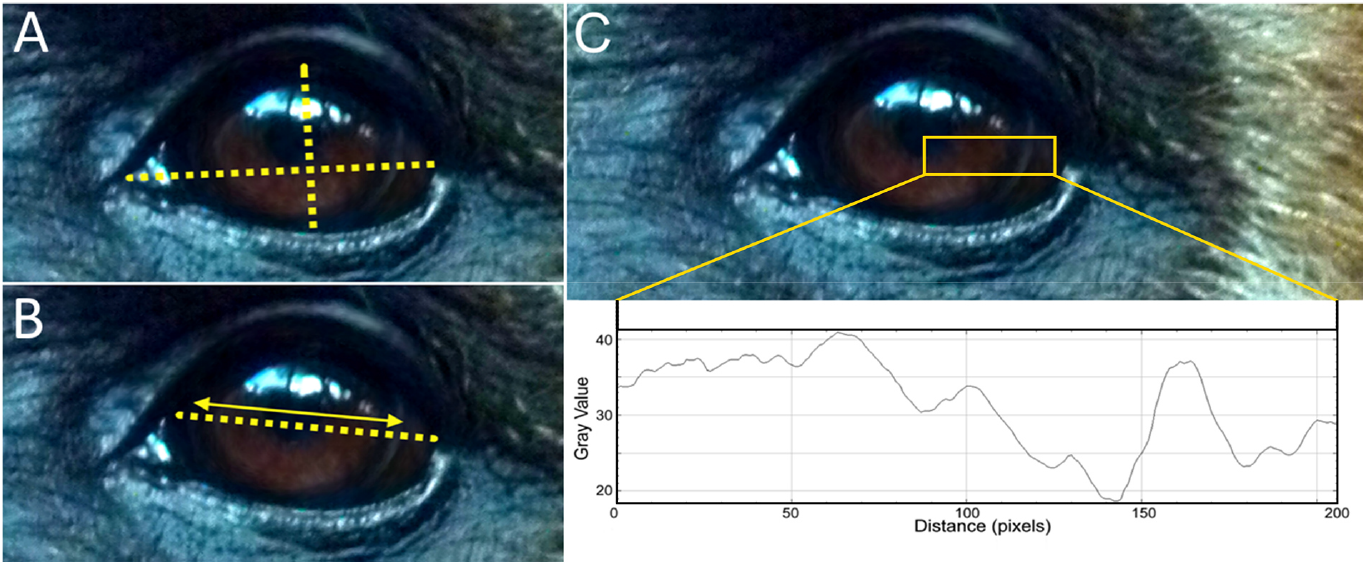
Visualization of measurements exemplified on a white-handed gibbon (*Hylobates lar*). A: Measurements of width and height of the eye taken to calculate a ratio (width-height ratio, WHR). B: Measurements of the width of visible sclera and iris to calculate a ratio (scleral size index, SSI). C: Deriving ocular luminance measurements (HC, RIL) from grey values in ImageJ using the *plot profile* function. Photo credit: Thomas Geissmann.

We used the *plot profile* function in ImageJ (Fig. 1) to retrieve luminance values from the images (Perea García, 2016). Only traits from one eye per subject were quantified. In doing so, we chose the better illuminated eye for measurements. In case both eyes were equally well visible (direct gaze conditions), we selected the one with higher contrast values.

We quantified grey scale luminance values to quantify ocular contrasts and classified eyes into two phenotypic groups following Perea García et al. (2019). In type 1 eyes, the sclera is lighter than the iris (e.g., humans), while the opposite is true for type 2 eyes (e.g., most chimpanzees). Dependent on eye type, we chose either the highest or lowest grayscale luminance values from either portion of the eye to achieve the highest possible contrast difference. Subsequently, these values were used to calculate the absolute (highest ocular contrast = HC) and relative (relative iris luminance = RIL) differences between scleral and iridal luminance per eye. RIL reflects the percentage of grayscale luminance shown by the darker portion of the eye (sclera or iris) in comparison to the lighter one, which per definition is assumed to represent 100% luminance (Perea García et al., 2017). Higher HC values indicate greater conspicuousness, while the opposite is the case for RIL. Procedures were adopted from Perea García (2016) and Perea García et al. (2019). Reflections mirrored in the eye as well as the demarcation line between iris and sclera were carefully avoided. We also quantified the grey value slope between iris and sclera (Perea García, 2016) for each individual but did not incorporate these data into our analyses (Suppl. Tab. 1).

Besides interspecific comparisons, we attempted to compare pigmentation patterns in wild and captive-bred individuals. However, sufficient samples for such a comparison could only be compiled for *Gorilla gorilla* (n_captive-bred_ = 23; n_wildborn_ = 17) and *Hylobates lar* (n_captive-bred_ = 13; n_wildborn_ = 9).

### Quantifying ocular shape and sclera exposure in hominoids

Ocular shape was quantified on the genus level for all genera of extant hominoids except for *Hoolock*. The latter was omitted due to a lack of suitable photographs. A set of images from 20 individuals representing either direct (eyes oriented towards the camera; n = 10) or sideways averted glance (n = 10) was analyzed for each genus, respectively. Congeneric species were grouped since preliminary screenings did not reveal notable differences. This resulted in a total sample of 140 pictures (Suppl. Tab. 2). In all these photos, subjects consistently faced the camera with fully opened eyes to reduce effects of the angle of photography on the measurements. While pictures of non-human primates were collected as described beforehand, all photos of humans derived from the private archives of the authors. All persons pictured gave consent for the photos to be used in this study. The human subjects were of diverse Eurasian descent. For nonhuman species, a trained observer (KRC) assigned photos into the “direct glance” and “averted glance” categories but a naive one (SB) scored them into said categories as well, allowing us to compute Cohen’s Kappa as a measure of inter-rater reliability.

The width-height ratio (WHR) and the exposed sclera size index (SSI) of eyes were quantified, following the procedures of Kobayashi and Kohshima (2001) and Mayhew and Gómez (2015) to allow for meaningful comparisons (Fig. 1). WHR is a measure to approximate eye shape, while SSI indicates the amount of exposed sclera. WHR was calculated from all images in the dataset for the respective genus, while SSI was calculated for direct and averted glance images independently to approximate changes in visible scleral surface during averted glancing. In case both eyes of an individual were clearly visible (n = 137), measurements from the left and right eye were averaged (Suppl. Tab. 2).

### Statistical analysis

All statistics were performed in R (R Core Team, 2020). After log-transformation, normal distribution of data was assessed by applying the Shapiro-Wilk test, homogeneity of variances by running the Bartlett test. Parametric data were compared via ANOVA, while non-parametric data were analyzed using Wilcoxon’s rank sum test. Tukey honest significant differences were employed as a post hoc test correcting for multiple comparisons subsequent to ANOVA, while Bonferroni correction was applied to address this issue for Wilcoxon tests. Differences in RIL and HC between wildborn and captive-bred individuals were tested by employing the Welch two sample t-test (*Hy. lar*) or the Wilcoxon rank sum test (*G. gorilla*). All tests were performed at a post-correction α-level of 0.05.

We visualized phylogenetic patterns, quantified phylogenetic signals and computed maximum likelihood ancestral state estimates with the *phytools* package version 0.7-70 (Revell, 2012). The hominoid phylogeny used was derived from the 10kTrees website (Arnold et al., 2010).

To visualize differences in the quantified ocular traits, a principal component analysis (PCA) was run on the species mean values for HC, RIL, and the species-specific proportion of type 1 eyes in the population (Type) as well as on the genus medians for SSI during averted glancing and WHR (see Tabs. 1 and 2). Due to the lack of SSI and WHR measurements, *Hoolock* was omitted from the analysis. SSI and WHR data of congeneric species were assumed to be equal.

**Table 2:** Median values (± SD) of width-height ratio (WHR) and sclera size index (SSI) during direct and averted gaze in seven hominoid genera.

| Genus | WHR | n | SSI (direct) | n | SSI (averted) | n |
| --- | --- | --- | --- | --- | --- | --- |
| *Hylobates* | 1.60 (0.13) | 20 | 1.15 (0.06) | 10 | 1.46 (0.13) | 10 |
| *Nomascus* | 1.57 (0.14) | 20 | 1.19 (0.09) | 10 | 1.36 (0.08) | 10 |
| *Symphalangus* | 1.70 (0.17) | 20 | 1.24 (0.06) | 10 | 1.46 (0.07) | 10 |
| *Gorilla* | 2.17 (0.24) | 20 | 1.71 (0.08) | 10 | 2.21 (1.30) | 10 |
| *Homo* | 2.75 (0.43) | 20 | 2.0 (0.31) | 10 | 2.30 (0.27) | 10 |
| *Pan* | 1.93 (0.16) | 20 | 1.46 (0.11) | 10 | 1.82 (0.16) | 10 |
| *Pongo* | 1.87 (0.15) | 20 | 1.52 (0.10) | 10 | 1.76 (0.28) | 10 |

## Results

### Qualitative assessment of ocular pigmentation in great and small apes

In small apes, the sclera was found to be predominately darker than the iris (type 2 phenotype; n = 135 of 168; 80 %), but notable interspecific variation was found (Tab. 1). In some species, this pattern was recovered for all individuals (*Ho. leuconedys*, *Hy. moloch*, *Hy. muelleri*, *N. leucogenys*). The highest prevalence of light sclerae (type 1 phenotype) was found in siamangs (*S. syndactylus*, n = 11 of 29, 38 %) and white-handed gibbons (*Hy. lar*, n = 14 of 30, 46 %). In most of these individuals, however, scleral depigmentation was moderate, leading to a medium to light brown sclera, which were often only minimally lighter than the iris. Of all sampled hylobatids, only four white-handed gibbons displayed advanced depigmentation bilateral of the sclera, resulting in a mottled white appearance, vaguely resembling that of humans. The small ape sclera, if not depigmented, appears dark brown to almost black in all species. Iris color was found to be mostly dark to chestnut brown (see Fig. 2). However, in *Hy. moloch* and *Hy. muelleri*, the iris has a dark amber color, similar to that of chimpanzees (Fig. 2).

**Figure 2:**
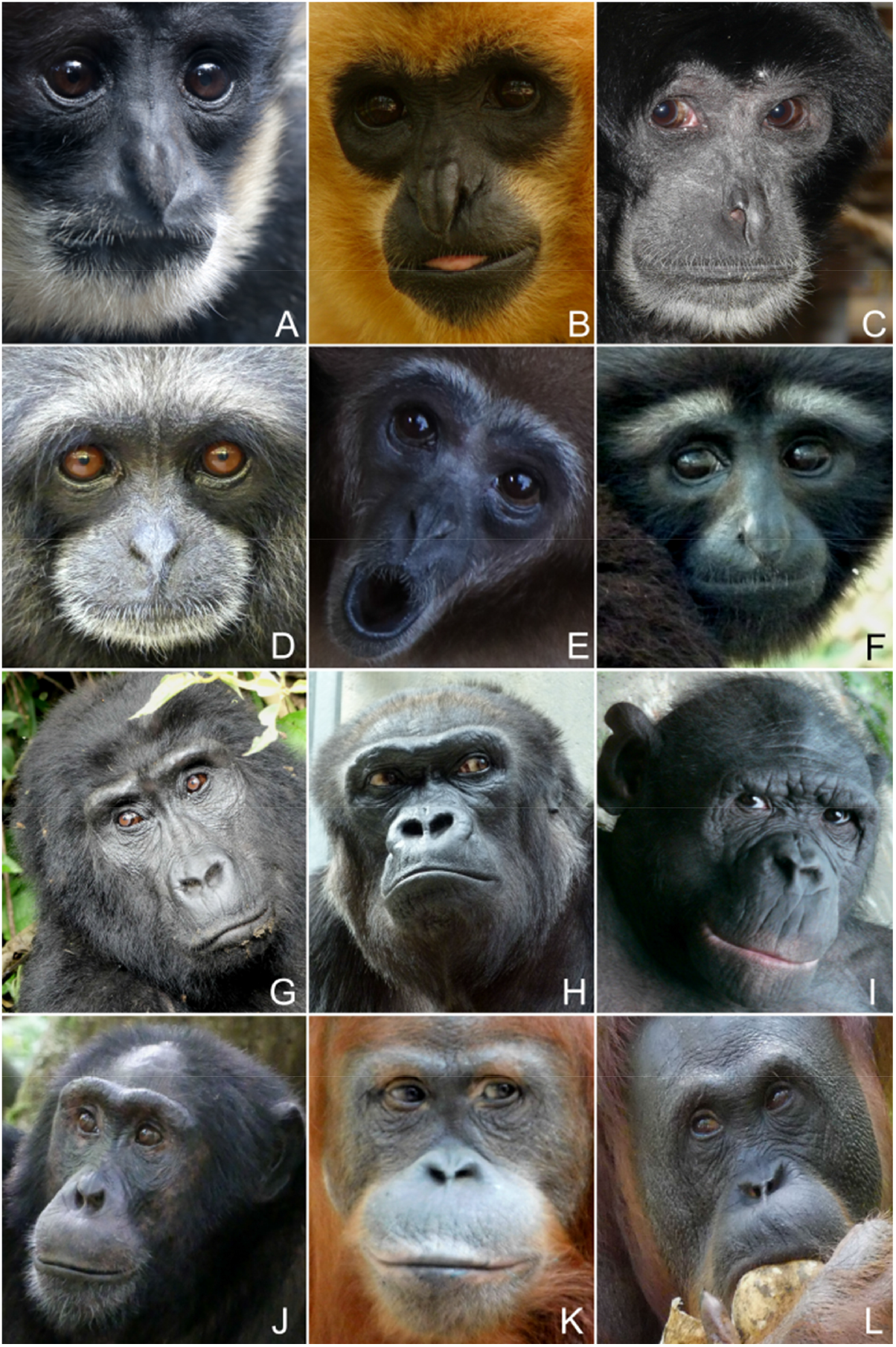
Typical external appearance of the eye in representatives of the seven non-human ape genera (superfamily Hominoidea). Note the difference in scleral exposure between gibbons (A-F) and great apes (G-L) A: Southern white-cheeked gibbon (*Nomascus (leucogenys) siki*). B: Southern yellow-cheeked gibbon (*Nomascus gabriellae*). C: Siamang (*Symphalangus syndactylus*). D: East Bornean gray gibbon (*Hylobates funereus*). E:White-handed gibbon (*Hylobates lar*). F: Gaoligong hoolock gibbon (*Hoolock tianxing*). G: Eastern gorilla (*Gorilla beringei*). H: Western gorilla (*Gorilla gorilla*). I: Bonobo (*Pan paniscus*). J: Chimpanzee (*Pan troglodytes*). K: Sumatran orangutan (*Pongo abelii*). L: Bornean orangutan (*Pongo pygmaeus*). Photo credit: A, E – Miriam Lindenmeier; K – Kai R. Caspar; all remaining pictures - Thomas Geissmann.

The sclerae of gorillas and orangutans tended to be at least in parts lighter than irises, displaying various degrees of depigmentation. Eastern gorillas were an exception to this trend. Here the inverse pattern (*G. beringei*, n = 13 of 22, 59 %) was found to be more abundant but both occur at high frequencies. In Western gorillas and orangutans light sclerae were predominately found (*G. gorilla*, n = 34 of 40, 85 %; *P. abelii*, n = 20 of 23, 87 %; *P. pygmaeus*, n = 19 of 24, 79 %). However, just as in many gibbon species, depigmentation in Bornean orangutans was often weak, leading to predominately brownish instead of white sclerae in this species. In both gorillas and orangutans, the portions of the sclera immediately surrounding the iris typically (but not always) remained pigmented, creating a dark frame of variable thickness.

Gorillas deviated from both orangutans and gibbons in frequently displaying clearly asymmetric depigmentation patterns. In approximately one quarter of Eastern (n = 6 of 22, 27 %) and Western gorillas (n = 10 of 40; 25 %), conspicuous depigmentation was restricted to just one eye. This pattern was only noted for one Sumatran orangutan and was absent in the Bornean species. In gibbons, such asymmetries were also rare, occurring in white-handed gibbons (n = 4 of 30, 13 %), pileated gibbons (n = 2 of 17, 12 %) and siamangs (n = 2 of 29, 7 %).

### Quantitative assessment of ocular pigmentation in great and small apes

On family level, we found a higher HC (Wilcoxon test: p << 0.001; Fig. 3) but lower RIL (Wilcoxon test: p << 0.001; Fig. 4) in hominids compared to hylobatids.

**Figure 3:**
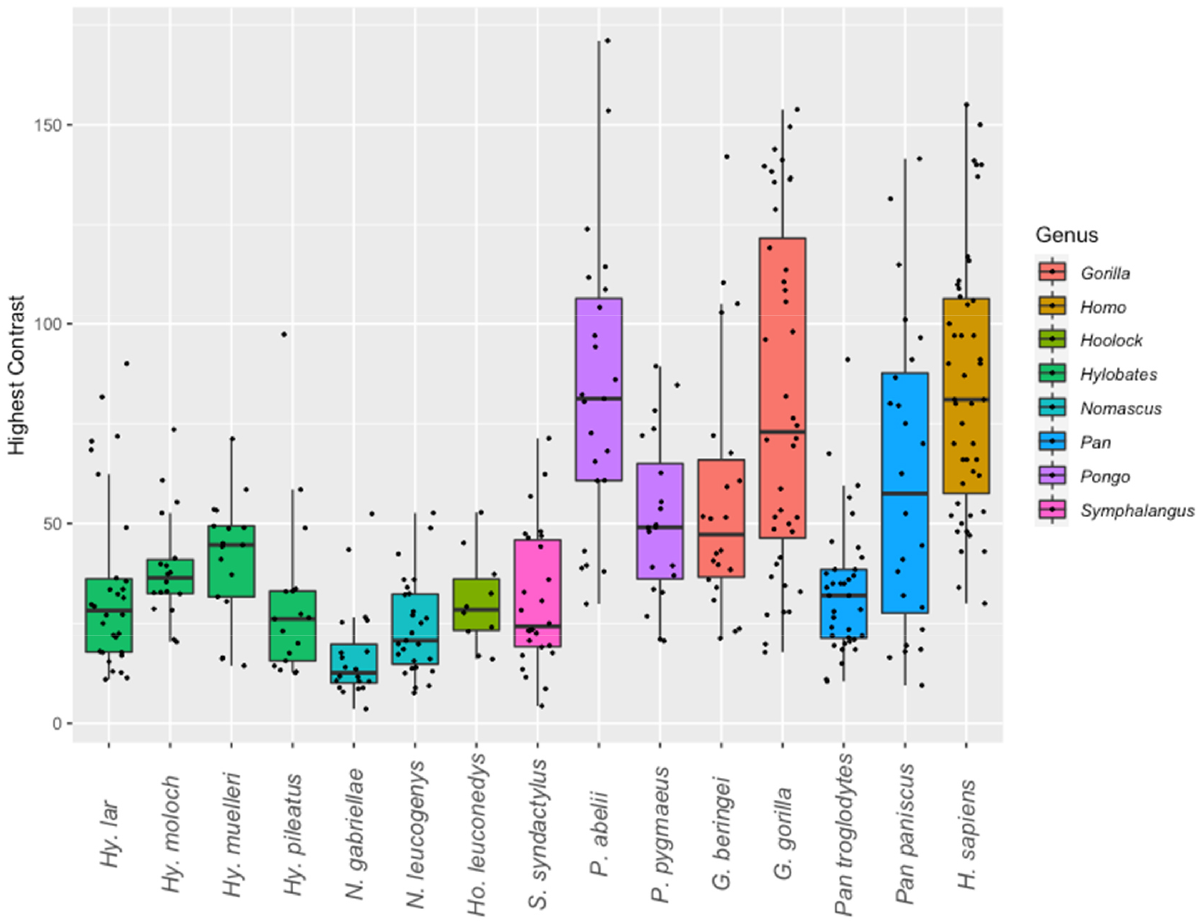
Variation of highest ocular contrast (HC) in the eight extant hominoid genera on species level.

**Figure 4:**
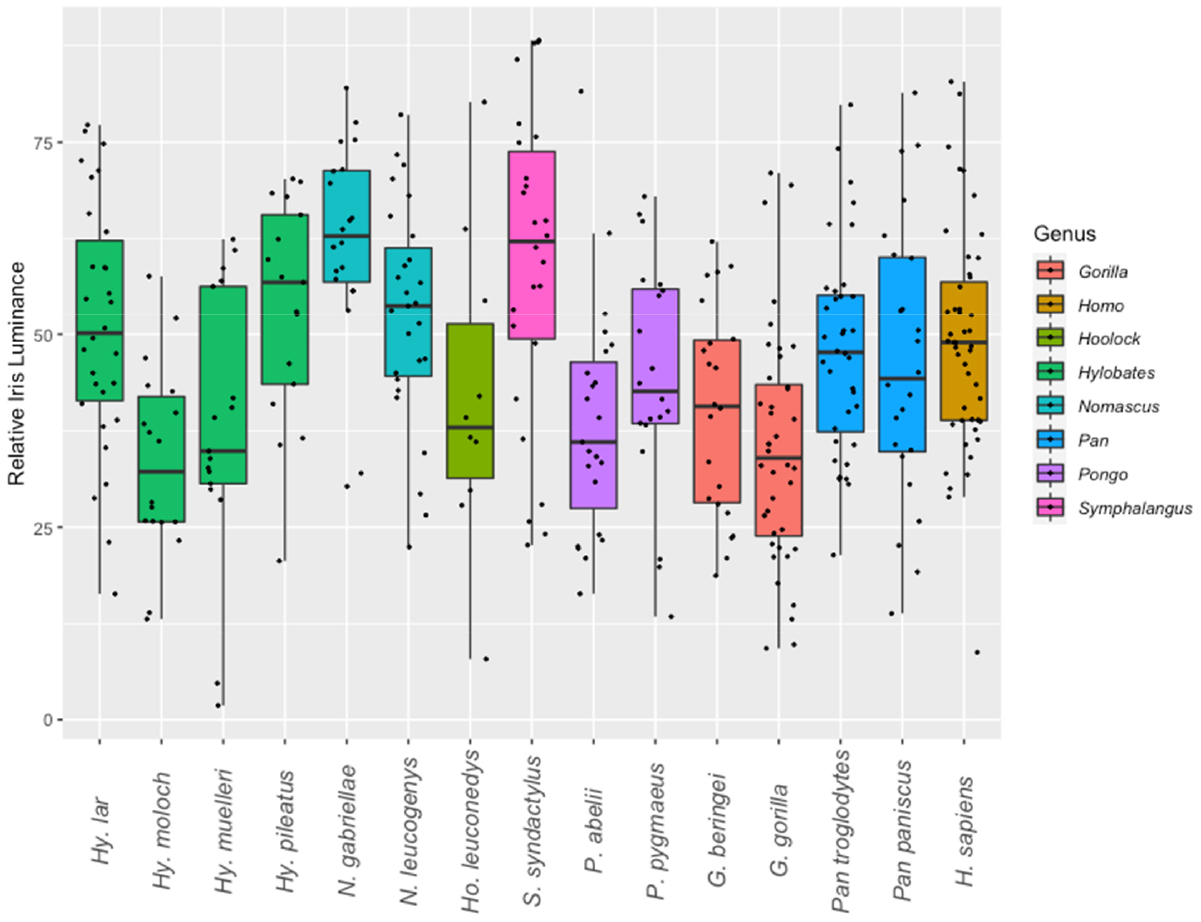
Variation of relative iris luminance (RIL) in the eight extant hominoid genera on species level.

Humans, Sumatran orangutans and Western gorillas exhibited the highest mean HC values, followed by bonobos, Eastern gorillas, Bornean orangutans and chimpanzees (Tab. 2). Chimpanzees were the only hominids for which the mean HC was recovered to lay within the range of variation of hylobatid species means. Among small apes, Bornean and Javan gibbons exhibited the highest HC values while the lowest were found among crested gibbons. Concerning RIL, species means for hominids fell within the range of hylobatid variation (Fig. 4). The lowest RIL in the sample was found for Javan gibbons, while the highest was recovered for Southern yellow-cheeked gibbons. Among hominids, humans displayed the highest RIL, while the lowest was found in Western gorillas (Tab. 2).

Reflecting these findings, we found a significant but moderate phylogenetic signal for HC among the hominoid sample (Pagel’s λ = 0.565, p = 0.026). RIL on the other hand was not found to correlate with phylogeny (Pagel’s λ << 0.001, p = 1), which is mirrored by the inconsistent distribution of the trait among the studied taxa. Maximum likelihood ancestral state estimates of HC and RIL for each node within our hominoid phylogeny are provided in Supplementary Table 3 and are visualized in Figure 5. Importantly, hominids, when compared to hylobatids, maintained a contrasting coloration throughout their evolutionary history. A comparatively high HC (63.6; 95% CI: 44-83.3) was also estimated for the common ancestor of the *Pan-Homo* clade, suggesting the dark eyes of chimpanzees to be a recently evolved trait.

**Figure 5:**
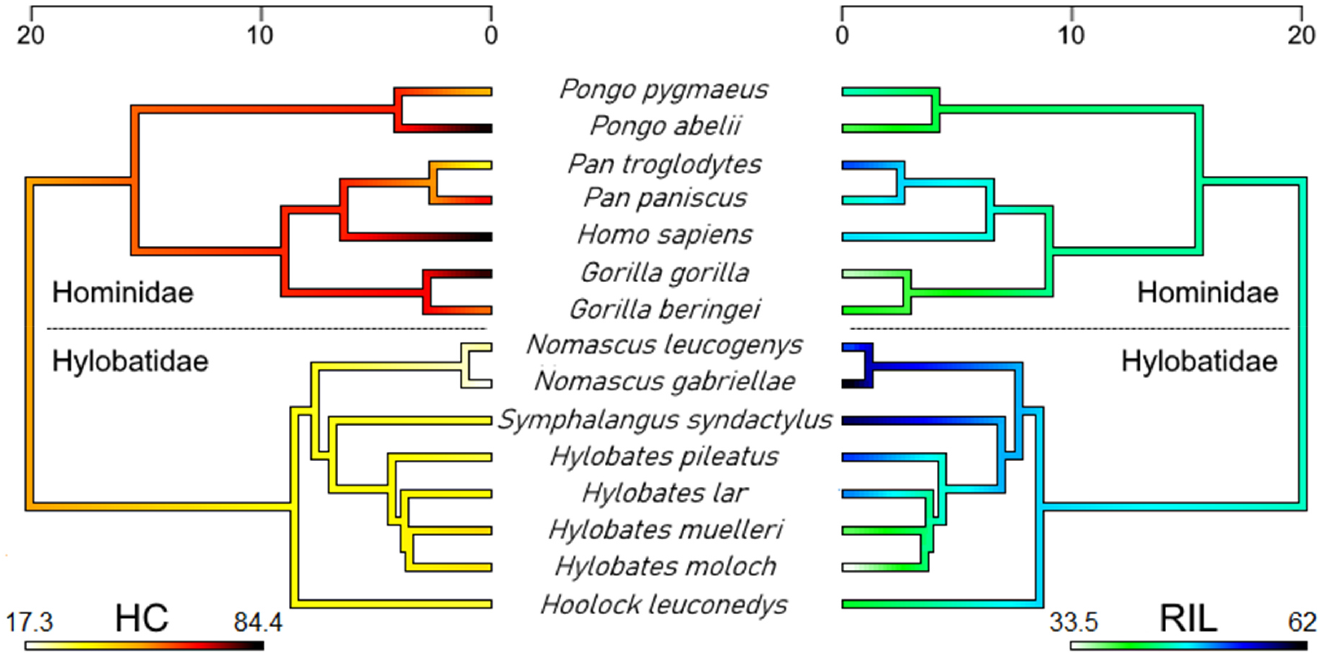
Visualization of phylogenetic patterns in species means of highest ocular contrast (HC, left) and relative iris luminance (RIL, right) in the ape superfamily (Hominoidea). Note the discrepancy between the two measures and the secondary acquisition of a gibbon-like HC in chimpanzees (*Pan troglodytes*). Time scales correspond to million years before present. Maximum likelihood ancestral state estimates at the nodes of the trees are provided in Supplementary Table 3.

Results from interspecies comparisons in HC and RIL are summarized in Table 3. For HC, only crested gibbons (genus *Nomascus*) exhibited values that significantly differed from other hylobatids, their eyes being notably dark. This genus also included the only gibbon species that significantly deviated from the patterns found in chimpanzees, bonobos and Bornean orangutans. Human HC differed significantly from all species in the sample except for Sumatran orangutans, bonobos, and gorillas, species which also tend to exhibit depigmented sclerae. Therefore, the contrasting of the human eye is not unique. Comparisons of RIL did not produce similarly comprehensible patterns. While some species did not show significant differences to any others in the sample (Eastern hoolock, bonobo), Western gorillas did so in comparison to seven species, including humans, chimpanzees, and a range of small apes. There was a moderate but significant negative correlation between HC and RIL (Pearson’s r = −0.54, p = 0.04).

**Table 3:** Species-level comparison of highest ocular contrast (HC) and relative iris luminance (RIL) between hominoids. Values of the upper triangular matrix (orange) correspond to statistical results for HC, those of the lower one (blue) to RIL. Bold values indicate significant results (HC: Bonferroni-corrected Wilcoxon tests; RIL: Tukey honest significant differences).

| Species | *Hoolock leuconedys* | *Hylobates lar* | *Hylobates moloch* | *Hylobates muelleri* | *Hylobates pileatus* | *Symphalangus syndactylus* | *Nomascus gabriellae* | *Nomascus leucogenys* | *Gorilla beringei* | *Gorilla gorilla* | *Homo sapiens* | *Pan paniscus* | *Pan troglodytes* | *Pongo abelii* | *Pongo pygmaeus* |
| --- | --- | --- | --- | --- | --- | --- | --- | --- | --- | --- | --- | --- | --- | --- | --- |
| *Hoolock leuconedys* |  | 1.000 | 1.000 | 1.000 | 1.000 | 1.000 | 0.289 | 1.000 | 0.973 | **0.01** | **0.001** | 1.000 | 1.000 | **0.001** | 0.648 |
| *Hylobates lar* | 0.942 |  | 1.000 | 1.000 | 1.000 | 1.000 | **0.018** | 1.000 | 0.129 | **0.000** | **0.000** | 0.755 | 1.000 | **0.000** | 0.355 |
| *Hylobates moloch* | 0.99 | **0.012** |  | 1.000 | 1.000 | 1.000 | **0.000** | **0.039** | 1.000 | **0.05** | **0.000** | 1.000 | 1.000 | **0.000** | 1.000 |
| *Hylobates muelleri* | 1.000 | 0.249 | 1.000 |  | 1.000 | 1.000 | **0.002** | 0.071 | 1.000 | 0.129 | **0.000** | 1.000 | 1.000 | **0.012** | 1.000 |
| *Hylobates pileatus* | 0.859 | 1.000 | **0.014** | 0.2 |  | 1.000 | 0.457 | 1.000 | **0.039** | **0.000** | **0.000** | 0.656 | 1.000 | **0.000** | 0.102 |
| *Symphalangus syndactylus* | 0.138 | 0.811 | **0.000** | **0.001** | 0.996 |  | 0.243 | 1.000 | 0.113 | **0.000** | **0.000** | 0.527 | 1.000 | **0.000** | 0.093 |
| *Nomascus gabriellae* | 0.055 | 0.49 | **0.000** | **0.000** | 0.934 | 1.000 |  | 1.000 | **0.000** | **0.000** | **0.000** | **0.001** | **0.005** | **0.000** | **0.000** |
| *Nomascus leucogenys* | 0.852 | 1.000 | **0.005** | 0.137 | 1.000 | 0.962 | 0.761 |  | **0.001** | **0.000** | **0.000** | **0.025** | 1.000 | **0.000** | **0.001** |
| *Gorilla beringei* | 1.000 | 0.424 | 0.99 | 1.000 | 0.345 | 0.002 | **0.001** | 0.25 |  | 1.000 | 0.074 | 1 | 0.073 | 0.886 | 1.000 |
| *Gorilla gorilla* | 0.997 | **0.002** | 1.000 | 1.000 | **0.005** | **0.000** | **0.000** | **0.001** | 0.996 |  | 1.000 | 1 | **0.000** | 1.000 | 1.000 |
| *Homo sapiens* | 0.99 | 1.000 | **0.024** | 0.417 | 1.000 | 0.302 | 0.113 | 1.000 | 0.643 | **0.003** |  | 1 | **0.000** | 1.000 | **0.01** |
| *Pan paniscus* | 1.000 | 0.999 | 0.326 | 0.928 | 0.984 | 0.175 | 0.063 | 0.983 | 0.99 | 0.244 | 1.000 |  | 1.000 | 1.000 | 1.000 |
| *Pan troglodytes* | 0.998 | 1.000 | 0.081 | 0.663 | 0.998 | 0.227 | 0.082 | 0.998 | 0.866 | **0.026** | 1.000 | 1.000 |  | **0.000** | 0.145 |
| *Pongo abelii* | 1.000 | 0.19 | 0.999 | 1.000 | 0.165 | **0.000** | **0.000** | 0.095 | 1.000 | 1.000 | 0.333 | 0.926 | 0.61 |  | 0.15 |
| *Pongo pygmaeus* | 1.000 | 0.973 | 0.687 | 0.996 | 0.911 | 0.081 | **0.027** | 0.894 | 1.000 | 0.677 | 0.998 | 1.000 | 1.000 | 0.997 |  |

No significant differences in RIL and HC could be detected between Western gorillas (Wilcoxon test: W ≥ 129; p ≥ 0.07) as well as white-handed gibbons (t-test: t ≥ −0.511; p ≥ 0.62) born in the wild compared to those bred in captivity.

### Quantifying ocular shape and sclera exposure in hominoids

Inter-rater reliability for the scoring of glance direction was strong throughout, but agreement was slightly higher for the great ape (59/60; κ = 0.96, p << 0.001) than for the gibbon sample (56/60; κ = 0.86; p << 0.001). Mean sclera size indices (SSI) were found to be consistently smaller in hylobatids compared to those of hominids, with a particularly pronounced difference occurring in the averted gaze condition (Tab. 2, Suppl. Tab. 4; Bonferroni-corrected Wilcoxon test: p < 0.001 for all hominid-hylobatid species pairs in both glance conditions). For width-heights-ratio (WHR), *Symphalangus* did not differ significantly from neither *Pan* nor *Pongo* (p > 0.13), while all other interfamilial comparisons of WHR yielded marked differences (p < 0.001). Small ape genera did not deviate significantly from each other in either direct or averted gaze SSI (p > 0.3) and the only significant difference in WHR among small apes was found between *Symphalangus* and *Nomascus* (p = 0.04), with the latter displaying lower WHR values. Within the Hominidae, *Homo* deviated significantly from all other genera in direct glance SSI and WHR (p ≤ 0.01). However, in averted glance SSI, *Homo* only differed significantly from *Pan* and *Pongo* (p < 0.05) but not from *Gorilla* (p = 1). No notable differences were found between *Pan* and *Pongo* in any of the measurements observed (p = 1 for all comparisons). *Gorilla* significantly deviated from all other hominids in WHR and direct glance SSI, but only from *Pan* regarding averted SSI (p < 0.01).

### Principal component analysis of ocular traits

PCA grouped hominoids into three groups based on ocular traits (Fig. 6). The first two principal components of the PCA encompassed 89.8 % of the total variance in the sample, exhibiting eigenvalues of 3.55 and 0.94, respectively. Variable contributions are visualized in Supplementary Figure 1. The first group is constituted by hylobatids and chimpanzees, which clustered together in the PCA morphospace (Fig. 6A). Bonobos grouped together with gorillas and orangutans, forming a second cluster. Finally, humans fell far outside of the range of variance of all the other hominoid genera. A second PCA omitting RIL, which is a problematic variable (see Discussion) resulted in a similar pattern (Suppl. Figs. 2 and 3), the first and second principal component encompassing even more of the total variance in the sample (94.3%). In this PCA, gorillas were situated closer to humans than in the first one, diffusing the cluster constituted by bonobos, gorillas and orangutans.

**Figure 6:**
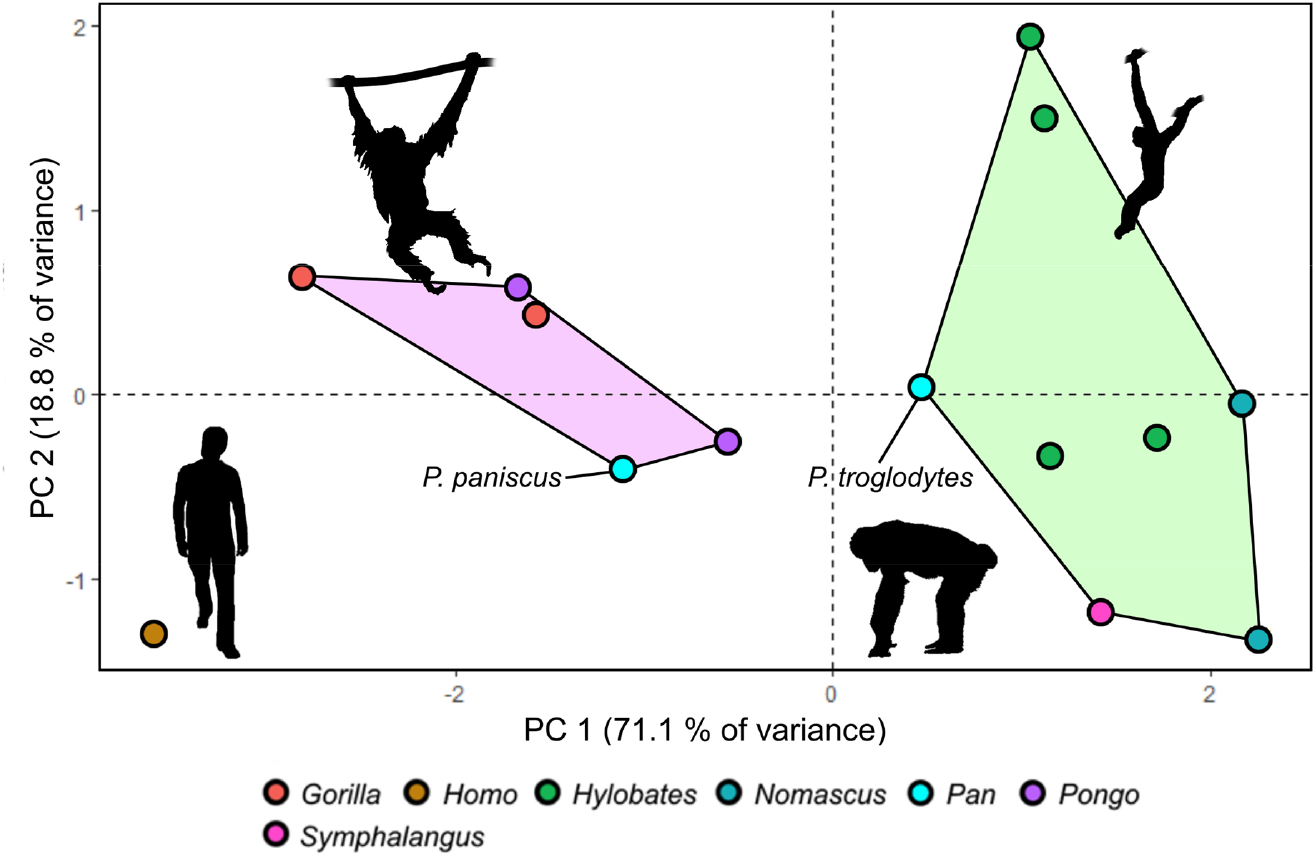
Results of PCA on quantified ocular traits in apes. A: Visualization of PCA results. Chimpanzees and gibbons form a cluster highlighted in green. Gorillas, bonobos and orangutans cluster together in a morphospace colored in light purple. Humans are separated from all other hominoids. The orangutan silhouette is in public domain, others were created by KRC.

## Discussion

### General discussion of results

Although we found only few consistent differences in ocular traits separating small and great apes, some predictions of the cooperative eye-hypothesis could be confirmed. Importantly, sclera size indices (SSI) were found to be consistently lower in gibbons compared to other apes and this difference became even more pronounced in averted gaze situations. This finding supports the conclusions of Kobayashi and Kohshima (2001) that were drawn from a small-scale dataset and further demonstrates that gibbon eyes are indeed far less suited to convey glance signals than those of great apes and humans. Regarding the width-height ratio of the eyes, hylobatids also displayed lower mean values than hominids. Nevertheless, siamangs approach the great ape genera *Pan* and *Pongo* in WHR and were found not to differ significantly from them in that regard. Still, this did not result in notably higher SSI values in siamangs compared to other gibbons, nor to equal SSI when compared to these great apes (Tab. 2). Our results on scleral exposure and width-height ratios in human and great ape eyes match the data from previous studies (Kobayashi & Kohshima, 2001; Kaplan & Rogers, 2002). In particular, we replicated the finding of Mayhew and Gómez (2015) that humans and gorillas do not differ in the degree of scleral exposure during averted glancing but do so in the direct glance condition. The reason for this disparity lies probably in the horizontally widened outline of the human eye. It causes rotations of the human eyeball to have less of an effect on scleral exposure when compared to other hominoids. Accordingly, the relative difference in the amount of visible sclera between direct and averted glance SSI shown by humans is exceeded by all great apes as well as by the gibbon genus *Hylobates* (Tab. 2).

Pigmentation patterns followed a roughly similar pattern to SSI among hominoids, but they were not congruent. Hylobatids displayed less contrasted eyes than their large-bodied relatives, as indicated by values for highest ocular contrast (HC), which were found to moderately correlate with hominoid phylogeny. As with the comparatively small amounts of exposed sclera in the gibbon eye, this again demonstrates a greater signaling value of hominid compared to hylobatid eyes. On the species level however, this notion cannot be generalized as chimpanzees were found to exhibit mean HC values in the range of gibbons, rendering their eyes similarly inconspicuous. Chimpanzee HC differed significantly from that of hominids with strongly contrasted eyes (humans, Sumatran orangutans, and Western gorillas) but not from the ones of siamangs, dwarf gibbons and hoolocks. As exemplified by our results on Western gorillas and white-handed gibbons, wildborn and captive-bred apes do not differ in ocular contrasts, making biases through imbalanced sampling of natural and captive populations unlikely. Still, it should be pointed out that our methodology does not capture the full extent of scleral pigmentation patterning. For instance, pronounced local scleral depigmentation will yield similar results to a fully depigmented sclera, despite pronounced phenotypic differences and related effects on glance direction signaling (compare Fig. 2C). This constitutes an important limitation of our method, which is also insensitive to asymmetric expressions of pigmentation. Merging quantitative analyses with a qualitative scoring of pigmentation patterns (compare Mayhew & Gómez, 2015) could constitute a way of overcoming this limitation in future studies. Another potential shortcoming of our, as well as of all previous approaches so far, is that only differences in ocular contrasts but not in hues are quantified and scored to approximate salience. This could have led to an underestimation of conspicuousness, particularly in species with dark sclerae.

Relative iris luminance (RIL) was recovered as a trait that varied independent of phylogeny. Importantly, we could not find support for the assumption that low RILs are reliable indicators of more conspicuously colored eyes (Perea García et al., 2019), despite a moderate negative correlation of the two traits within our sample. The lowest average RIL value found was that of the Javan gibbon (*Hylobates moloch*, mean RIL: 33.5) which shows an amber-colored iris and dark brown sclera, rendering its eyes obviously far less conspicuous than the ones of, for instance, humans (mean RIL: 48.7) which were found to have a significantly higher RIL. This fact points out a major issue in the usage of RIL as a meaningful measure of ocular pigmentation. If gray values for sclera and iris are both low and differ little from one another, resulting RIL values may still equal or range below those obtained from eyes that show salient contrasts between these regions. This insensitivity makes RIL an unsuitable proxy for the conspicuousness of ocular pigmentation patterns. However, it might still be used to analyze intraspecific pigmentation patterns in groups with uniformly colored sclerae such as humans, for which the measure has originally been established (Perea García et al., 2017). Given that HC is a more faithful measure of general ocular conspicuousness, we propose to rely on HC rather than on RIL to quantify ocular pigmentation in future comparative studies. For this, large sample sizes are recommended to counteract effects of differing lighting regimes in the analyzed pictures. (Perea García et al., 2019) have argued based on RIL, that chimpanzee, bonobo and human eyes are equally suited to convey gaze signals. However, for reasons just pointed out, this argument does not hold. In ocular pigmentation, chimpanzees’ dark eyes resemble the ones of gibbons more than the human or even the average bonobo condition, arguing against a human-like social signaling function of chimpanzee eye coloration (compare Fig. 4). Above that, scleral exposure in *Pan* is the lowest among African apes, differing significantly from both humans and gorillas in averted and direct glancing situations. For these reasons, chimpanzee eyes are notable for being, on average, the least conspicuous of all hominids. Low RIL values alone fail to diagnose species that employ glance cueing or even sophisticated gaze cueing, as exemplified by several gibbon species in our sample.

Our results highlight differences in scleral depigmentation rates in hominids compared to hylobatids. The dataset of Kobayashi and Kohshima (2001) would suggest the dark-eyed gibbon pattern to be the plesiomorphic one. Therefore, more contrastingly colored eyes would be expected to have evolved in hominids after their split from hylobatids. However, this hypothesis can be challenged, given the small species sample sizes in that study together with the fact that its assumptions were later shown not to hold for most hominids (Mayhew & Gómez, 2015; Perea García, 2016; Perea García et al., 2019). From just our own experience, we can anecdotally report the presence of light sclerae in multiple species of Old World and New World monkeys (Suppl. Fig. 4). Studies on the frequency and phylogenetic distribution of this trait in primates other than apes are necessary to sufficiently characterize the ancestral state for hominoid eye pigmentation.

It is difficult to discern what underlies patterns of ocular appearance among the two hominoid families. Considering the *cooperative eye hypothesis*, it might be tempting to suggest that brighter and more exposed eyes in great apes reflect the more sophisticated gaze following behavior in this group when compared to gibbons (Liebal & Kaminski 2012). However, the hypothesis fails to explain the derived chimpanzee phenotype and cannot account for the great variability of hominid ocular contrasts. Furthermore, differences in SSI and WHR between hylobatids and hominids might be more parsimoniously explained by scaling effects deriving from differences in body size instead of by communicative demands (Kobayashi & Kohshima, 2001).

It is notable that each great ape genus encompasses species that markedly differ in their ocular pigmentation patterns (compare Perea García, 2016; Perea García et al., 2019). Perea García et al. (2019) suggested that scleral depigmentation in apes might be an evolutionary byproduct of greater social tolerance, induced by pleiotropic genes controlling neural crest development and mirroring patterns found in domesticated mammals. Indeed, the dark eyed Bornean orangutans and Eastern gorillas are less tolerant towards unfamiliar conspecifics than their congeners in the wild (van Schaik, 1999; Cooksey et al. 2020). Nevertheless, it remains to be clarified whether this is a consequence of intrinsic behavioral predispositions rather than of extrinsic factors relating to habitat characteristics. Between bonobos and chimpanzees, the former are far better characterized than in other ape taxa and point to marked physiological differences underlying behavioral disparities within the genus *Pan* (Wobber et al. 2010; Rilling et al. 2012;). Comparisons of socially tolerant and despotic monkey species could further test for a correlation between scleral pigmentation and aggressiveness. Yet, specific pleiotropic effects affecting scleral but not general skin or fur coloration appear to be yet undescribed in mammals (compare e.g., Cieslak et al., 2011), making the link a speculative one at the moment. It is also unclear, how the social tolerance hypothesis might apply to the hominid family in general when compared to hylobatids or specifically to white-handed gibbons. Although the latter show the highest scleral depigmentation rates among small apes, there is no evidence to suggest them to exhibit decreased levels of aggression towards unfamiliar conspecifics when compared to other species.

To sum up, the *cooperative eye hypothesis* might fit the family-level patterns we describe but loses its explanatory power on the species level. It is further important to point out that the human eye is on average not more saliently contrasted than that of gorillas, bonobos and Sumatran orangutans, further challenging its validity. The occurrence of at least locally depigmented sclerae in varying portions of the total population likely is an ancient hominid trait but its biological significance remains obscure. The view that extensive scleral depigmentation is exclusive to the human lineage (Kobayashi & Kohshima, 2001) or even diagnostic for the species *Homo sapiens* (Hare, 2017) cannot be maintained. Finally, reliance on RIL as an indicator of ocular conspicuousness may give rise to misleading results.

### Is ocular pigmentation linked to specific sociocognitive traits?

Previous research on scleral pigmentation in primates has highlighted a potential connection between cognitive traits such as social glance cueing and conspicuous ocular pigmentations (Kobayashi & Kohshima, 2001; Tomasello et al., 2007; Hare, 2017; Perea García et al., 2019). Whether these characters are correlated is not yet clear, however, since humans are the only primate species that evidently combines them (see below). Deducing glance cueing and other cognitive abilities from eye coloration alone may quickly lead into an adaptationist pitfall. Even if primate species converge in ocular pigmentation, the ways in which these species perceive conspecifics’ eyes may differ. For example, Western gorillas and Sumatran orangutans exhibit ocular contrasts resembling the human condition. Yet, their viewing patterns of conspecific’s faces and particularly eyes, is more reminiscent of chimpanzees than humans, they exhibit pronounced gaze avoidance in diverse social contexts and evidence of conspecific glance cue exploitation is, to our knowledge, absent (Kaplan & Rogers, 2002; Kano et al., 2012).

The hypothesis that dark eyes evolved to mask glance direction in competitive social environments must be critically reevaluated as well. Kobayashi and Kohshima (2001) proposed that all non-human primates would exhibit dark sclerae to benefit from “gaze camouflage”, but this idea remains a hypothetical rather than an empirical one. This assumption is not based on any experimental evidence and does not address how this trait would be advantageous to primates with low SSI and across the extreme diversity of social, activity and foraging regimes that the 91 species they studied encompass. In line with this, the additional notion that dark sclerae would lower predation risk via gaze concealment (Langton et al., 2000; Kobayashi & Kohshima, 2001) is also purely speculative. Kobayashi and Kohshima (2001) also ignore the relevance of other facial ornaments for gaze/glance communication (compare Kaplan & Rogers, 2002 and Ueda et al., 2014) and the great variety of cooperative behaviors in non-human primates that are often linked to gaze following (compare Micheletta & Waller, 2012). Dark sclerae appear to be a plesiomorphic primate trait (Kobayashi & Kohshima, 2001) and disentangling its evolutionary roots will require to take information on other mammals or even more distantly related vertebrate groups into account. We therefore discard both the glance cueing and cryptic gaze hypotheses of eye coloration as too simplistic to be helpful in interpreting the evolution of ocular pigmentation in primates. Our dataset could be expanded to test, whether ocular morphology indeed correlates with specific cognitive traits across primate groups. For this, however, there must be agreement on the definitions of the cognitive characteristics studied. At the moment, this is not the case for primate gaze or glance cueing.

So far, most information on gaze and glance cue understanding in apes derive from studies in which human experimenters signal to an ape subject in a captive setting (Barth et al., 2005; Tomasello et al., 2007; Caspar et al., 2018; Sanchez-Amaro et al., 2020). It is important to note that responsiveness to human eye orientation by habituated animals does not equate with the usage of glance cues among conspecifics, as for instance the successful exploitation of human glances by Californian sea lions (*Zalophus californianus*) demonstrates (Arkwright et al., 2016). The evidence, that apes utilize conspecifics’ glancing independent from head orientation to inform their actions is meager. Many studies assume that apes’ head direction and eye orientation align with each other for the most part, despite evidence of the contrary (Bethell et al., 2007). Equating head and eye direction in interpretations of glancing behavior has at times led to confusion. For example, Perea García et al. (2019) cite studies that either did not differentiate between the two (Kano & Call, 2014; Lucca et al., 2018) or that approximated gaze by head orientation alone (Hall et al., 2017) to support the notion that glance cues are relevant to chimpanzee communication. We are not aware of studies that unequivocally show exploitation of conspecific glance cues in any ape species within a social context. An effect of ocular contrast on such behaviors would need to be demonstrated further. Since humans can reliably deduce the glance direction of chimpanzees, it can be hypothesized that from a perceptual perspective, conspecifics might do so as well, despite their cryptically colored eyes (Bethell et al., 2007). The effect of eye color on gaze salience might well be minor. On a different note, it should be discussed how exactly the inclusion of glance cues could enhance apes’ communicative repertoire in the wild. What referential information could glance cues convey in a naturalistic setting that head orientation cannot? In the absence of evidence for conspecific glance cueing in great and small apes or an unambiguous link between ocular pigmentation and cognitive traits relating to gaze/glance following, as well as for reasons of parsimony, we assume that these characters evolve independently from one another.

### Which evolutionary factors underly the pigmentation of the human eye?

The evolutionary trend of scleral depigmentation in hominids finds its strongest expression in humans. Although rudimentary pigmentation of the conjunctiva and inconspicuous scleral spots can frequently be found in humans (Yanoff, 1969), apparent complete scleral depigmentation approaches 100 % in our species (Kobayashi & Kohshima, 2001). Why do humans, but not other apes, exhibit such a uniformly white scleral phenotype? The assumption that this trait evolved to facilitate glance or gaze cueing is problematic. First, as already pointed out, its occurrence among mammals that do or do not show sophisticated gaze following or glance cueing has not been sufficiently investigated. Above that, available evidence suggests that humans can reliably assess the glance direction of chimpanzees from a distance of 2–10 m (Bethell et al., 2007) as well as that of human models with artificially modified scleral colors (Yorzinski & Miller, 2020). Although it took naïve participants significantly longer to assess the glance direction of the latter compared to natural human eyes, the time differences only encompassed fractions of a second and no differences in the accuracy of deducing glance direction from normal human models and those with matched iris and scleral colors were found (Yorzinski & Miller, 2020). The relevance of the depigmented sclera for human communication might therefore be overstated, especially when compared to other morphological (i.e., the widened horizontal outline of the visible eyeball) or physiological ocular traits of our species (i.e., emotional tearing), which are not shared by other hominoids (Gračanin et al., 2018; Mayhew & Gómez, 2015; Fig. 4).

What requires an explanation is perhaps not that the human sclera simply exhibits depigmentation but that its expression is uniformly extreme across individuals and populations. The marked variability of scleral color in other hominid genera, which includes complete scleral depigmentation (Mayhew & Gómez 2015), makes a similar phenotypic diversity in human ancestors appear likely. Following that, the human pattern does not necessarily require an adaptive explanation but may simply result from genetic drift acting on ancestral trait variability. It is also reasonable to assume that sexual selection has contributed to the evolution of human eye pigmentation, not excluding but possibly complementing effects of genetic drift. As we have shown, other apes do not only exhibit strong interindividual differences in scleral pigmentation but at times also asymmetric scleral coloration, particularly gorillas (compare also Mayhew & Gómez (2015)). Both may point to a relaxed evolutionary pressure on ocular appearance in great apes compared to humans.

It has been demonstrated that scleral brightness strongly affects the attractiveness of human eyes and that it can act as an indicator of individual age (Gründl et al., 2012; Russell et al., 2014). A negative effect of age on scleral brightness has also been shown for chimpanzees and bonobos (Perea García et al., 2019). Thereby, uniformly light, salient sclerae should contribute to a juvenilized appearance of the face, which complies to general sexual selection pressures for neotenic facial traits in humans compared to other hominids, particularly in females (Jones et al., 1995; Perrett et al., 1998; Penin et al., 2002; Hare, 2017). Similarly, the symmetrical scleral depigmentation pattern in our species is in line with that human facial attractiveness is enhanced by increased symmetry (Thornhill & Gangestad, 1999). Light sclerae in humans could take over signaling functions that are absent in other apes, for example because they pay less attention to or even prefer to avoid gazing into each other’s eyes, e.g., gorillas and orangutans (Kaplan & Rogers, 2002; Kano et al., 2012); or because correlates of scleral brightness such as youthfulness do not increase sexual attractiveness in these species, e.g., chimpanzees (Muller et al., 2006). Distinct selection pressure on ocular appearance in humans is also indicated by iris color variability. Although there is at least one other primate genus, spider monkeys, in which two distinct iris color morphs co-occur in some species (*Ateles fusciceps* and *A. hybridus* - Meyer et al., 2013; *A. paniscus* - personal observation), humans appear to be unique among nondomesticated mammals in the diversity of iridal hues found in the global population, particularly Western Asia and Europe (Negro et al., 2017). Other apes display uniform, species-specific iridal coloration. Given that humans exhibit notable eye color preferences in the context of mate choice (Štěrbová et al., 2019), it can be assumed that iris color represents a sexually selected ocular trait that differentiates our species from non-human apes, just as it might be the case with the plain white sclera.

## Conclusion

Our data add to growing evidence suggesting a graded evolution of hominoid ocular coloration instead of a clear dichotomy between human and non-human primate eyes. Importantly, the evolutionary drivers of apparently derived scleral depigmentation in hominids, which peaked in humans and reversed in chimpanzees, remain unidentified. Still, the great intra- and interspecific variability opens possibilities for comparative research which should include both the great apes and gibbons as well as a range of outgroup taxa. The classic *cooperative eye hypothesis* that proposes an evolutionary link between primate ocular morphology and social signaling, needs to be experimentally revisited and scrutinized. In order to uphold it, a clear relevance to ocular pigmentation traits for communication among conspecifics in nonhuman primates needs to be demonstrated.

## Acknowledgements

KRC was supported by a PhD fellowship of the German National Academic Foundation (Studienstiftung des deutschen Volkes). We thank Miriam Lindenmeier for the permission to use selected photographs in this manuscript and Marie Padberg for helpful comments on the text. Finally, we acknowledge the many photographers who publicly shared their primate photos online, allowing us to analyze them for this study. The stunning ape pictures by Arjan Haverkamp proved to be of particular value for our study.

## References

Arkwright, T., Malassis, R., Carter, T., & Delfour, F. (2016). California sea lions (*Zalophus californianus*) can follow human finger points and glances. International Journal of Comparative Psychology, 29.

Arnold, C., Matthews, L. J., & Nunn, C. L. (2010). The 10kTrees website: A new online resource for primate phylogeny. Evolutionary Anthropology: Issues, News, and Reviews, 19(3), 114–118.

Barth, J., Reaux, J. E., & Povinelli, D. J. (2005). Chimpanzees’ (*Pan troglodytes*) use of gaze cues in object-choice tasks: different methods yield different results. Animal Cognition, 8(2), 84–92.

Bethell, E. J., Vick, S.-J., & Bard, K. A. (2007). Measurement of eye-gaze in chimpanzees (*Pan troglodytes*). American Journal of Primatology, 69(5), 562–575.

Burgin, C. J., Wilson, D. E., Mittermeier, R. A., Rylands, A. B., Lacher, T. E., & Sechrest, W. (2020). Illustrated Checklist of the Mammals of the World, Vol. 1: Monotremata to Rodentia. Barcelona: Lynx Editions.

Butler, D., & Suddendorf, T. (2014). Reducing the neural search space for hominid cognition: What distinguishes human and great ape brains from those of small apes? Psychonomic Bulletin & Review, 21(3), 590–619.

Caspar, K. R., Mader, L., Pallasdies, F., Lindenmeier, M., & Begall, S. (2018). Captive gibbons (Hylobatidae) use different referential cues in an object-choice task: insights into lesser ape cognition and manual laterality. PeerJ, 6, e5348.

Cieslak, M., Reissmann, M., Hofreiter, M., & Ludwig, A. (2011). Colours of domestication. Biological Reviews, 86(4), 885–899.

Deaner, R. O., & Platt, M. L. (2003). Reflexive social attention in monkeys and humans. Current Biology, 13(18), 1609–1613.

Geissmann, T. (2003). Circumfacial markings in siamang and evolution of the face ring in the Hylobatidae. International Journal of Primatology, 24(1), 143–158.

Goodall, J. (1986). The Chimpanzees of Gombe: Patterns of Behavior. Cambridge, MA: Belknap Press.

Gračanin, A., Bylsma, L. M., & Vingerhoets, A. J. J. M. (2018). Why only humans shed emotional tears. Human Nature, 29(2), 104–133.

Gründl, M., Knoll, S., Eisenmann-Klein, M., & Prantl, L. (2012). The blue-eyes stereotype: do eye color, pupil diameter, and scleral color affect attractiveness? Aesthetic Plastic Surgery, 36(2), 234–240.

Hall, K., Oram, M. W., Campbell, M. W., Eppley, T. M., Byrne, R. W., & de Waal, F. B. M. (2017). Chimpanzee uses manipulative gaze cues to conceal and reveal information to foraging competitor. American Journal of Primatology, 79(3), e22622.

Hare, B. (2017). Survival of the friendliest: *Homo sapiens* evolved via selection for prosociality. Annual Review of Psychology, 68(1), 155–186.

Jones, D., Brace, C. L., Jankowiak, W., Laland, K. N., Musselman, L. E., Langlois, J. H., … Symons, D. (1995). Sexual selection, physical attractiveness, and facial neoteny: cross-cultural evidence and implications. Current Anthropology, 36(5), 723–748.

Kano, F., & Call, J. (2014). Cross-species variation in gaze following and conspecific preference among great apes, human infants and adults. Animal Behaviour, 91, 137–150.

Kano, F., Call, J., & Tomonaga, M. (2012). Face and eye scanning in gorillas (*Gorilla gorilla*),orangutans (*Pongo abelii*), and humans (*Homo sapiens):* Unique eye-viewing patterns in humans among hominids. Journal of Comparative Psychology, 126(4), 388–398.

Kaplan, G., & Rogers, L. J. (2002). Patterns of gazing in orangutans (*Pongo pygmaeus*). International Journal of Primatology, 23(3), 501–526.

Kobayashi, H., & Kohshima, S. (2001). Unique morphology of the human eye and its adaptive meaning: comparative studies on external morphology of the primate eye. Journal of Human Evolution, 40(5), 419–435.

Langton, S. R. H., Watt, R. J., & Bruce, V. (2000). Do the eyes have it? Cues to the direction of social attention. Trends in Cognitive Sciences, 4(2), 50–59.

Liebal, K., & Kaminski, J. (2012). Gibbons (*Hylobates pileatus, H. moloch, H. lar, Symphalangus syndactylus*) follow human gaze, but do not take the visual perspective of others. Animal Cognition, 15(6), 1211–1216.

Lord, K. A., Larson, G., Coppinger, R. P., & Karlsson, E. K. (2020). The history of farm foxes undermines the animal domestication syndrome. Trends in Ecology & Evolution, 35(2), 125–136.

Lucca, K., MacLean, E. L., & Hare, B. (2018). The development and flexibility of gaze alternations in bonobos and chimpanzees. Developmental Science, 21(4), e12598.

ManyPrimates, Altschul, D. M., Beran, M. J., Bohn, M., Caspar, K. R., Fichtel, C., … Watzek, J. (2019). Collaborative open science as a way to reproducibility and new insights in primate cognition research. Japanese Psychological Review, 62(3), 205–220.

Mayhew, J. A., & Gómez, J.-C. (2015). Gorillas with white sclera: a naturally occurring variation in a morphological trait linked to social cognitive functions. American Journal of Primatology, 77(8), 869–877.

Meyer, W. K., Zhang, S., Hayakawa, S., Imai, H., & Przeworski, M. (2013). The convergent evolution of blue iris pigmentation in primates took distinct molecular paths. American Journal of Physical Anthropology, 151(3), 398–407.

Micheletta, J., & Waller, B. M. (2012). Friendship affects gaze following in a tolerant species of macaque, *Macaca nigra*. Animal Behaviour, 83(2), 459–467.

Morris, D. (1985). Bodywatching - A Field Guide to the Human Body. London: Jonathan Cape.

Muller, M. N., Thompson, M. E., & Wrangham, R. W. (2006). Male chimpanzees prefer mating with old females. Current Biology, 16(22), 2234–2238.

Negro, J. J., Carmen Blázquez, M., & Galván, I. (2017). Intraspecific eye color variability in birds and mammals: a recent evolutionary event exclusive to humans and domestic animals. Frontiers in Zoology, 14(1), 53.

Penin, X., Berge, C., & Baylac, M. (2002). Ontogenetic study of the skull in modern humans and the common chimpanzees: Neotenic hypothesis reconsidered with a tridimensional procrustes analysis. American Journal of Physical Anthropology, 118(1), 50–62.

Perea García, J. O. (2016). Quantifying ocular morphologies in extant primates for reliable interspecific comparisons. Journal of Language Evolution, 1(2), 151–158.

Perea García, J. O., Grenzner, T., Hešková, G., & Mitkidis, P. (2017). Not everything is blue or brown: quantification of ocular coloration in psychological research beyond dichotomous categorizations. Communicative & Integrative Biology, 10(1), e1264545.

Perea García, J. O., Kret, M. E., Monteiro, A., & Hobaiter, C. (2019). Scleral pigmentation leads to conspicuous, not cryptic, eye morphology in chimpanzees. Proceedings of the National Academy of Sciences, 116(39), 19248.

Perrett, D. I., Lee, K. J., Penton-Voak, I., Rowland, D., Yoshikawa, S., Burt, D. M., … Akamatsu, S. (1998). Effects of sexual dimorphism on facial attractiveness. Nature, 394(6696), 884–887.

Povinelli, D. J., & Eddy, T. J. (1996). Chimpanzees: joint visual attention. Psychological Science, 7(3), 129–135.

R Core Team (2020). R: A language and environment for statistical computing. Vienna, Austria: R Foundation for Statistical Computing.

Reichard, U. H., Barelli, C., Hirai, H., & Nowak, M. G. (2016). The evolution of gibbons and siamang. In U. H. Reichard, H. Hirai, & C. Barelli (Eds.), Evolution of Gibbons and Siamang: Phylogeny, Morphology, and Cognition (pp. 3–41). New York, NY: Springer New York.

Revell, L. J. (2012). phytools: an R package for phylogenetic comparative biology (and other things). Methods in Ecology and Evolution, 3(2), 217–223.

Russell, R., Sweda, J. R., Porcheron, A., & Mauger, E. (2014). Sclera color changes with age and is a cue for perceiving age, health, and beauty. Psychology and Aging, 29(3), 626–635.

Sanchez-Amaro, A., Tan, J., Kaufhold, S. P., & Rossano, F. (2020). Gibbons exploit information about what a competitor can see. Animal Cognition 23(2), 289–299.

Schneider, C. A., Rasband, W. S., & Eliceiri, K. W. (2012). NIH Image to ImageJ: 25 years of image analysis. Nature Methods, 9(7), 671–675.

Štěrbová, Z., Tureček, P., & Kleisner, K. (2019). Consistency of mate choice in eye and hair colour: testing possible mechanisms. Evolution and Human Behavior, 40(1), 74–81.

Thornhill, R., & Gangestad, S. W. (1999). Facial attractiveness. Trends in Cognitive Sciences, 3(12), 452–460.

Tomasello, M., Hare, B., Lehmann, H., & Call, J. (2007). Reliance on head versus eyes in the gaze following of great apes and human infants: the cooperative eye hypothesis. Journal of Human Evolution, 52(3), 314–320.

Ueda, S., Kumagai, G., Otaki, Y., Yamaguchi, S., & Kohshima, S. (2014). A comparison of facial color pattern and gazing behavior in canid species suggests gaze communication in gray wolves (*Canis lupus*). PLOS ONE, 9(6), e98217.

Wright, D., Henriksen, R., & Johnsson, M. (2020). Defining the domestication syndrome: comment on Lord et al. 2020. Trends in Ecology & Evolution, 35(12), 1059–1060.

Yanoff, M. (1969). Pigment spots of the sclera. Archives of Ophthalmology, 81(2), 151–154.

Yorzinski, J. L., & Miller, J. (2020). Sclera color enhances gaze perception in humans. PLOS ONE, 15(2), e0228275.

