## Supplementary Figures for "Ocular pigmentation in humans, great apes, and gibbons is not suggestive of communicative functions - having an eye on the ‘cooperative eye hypothesis’"

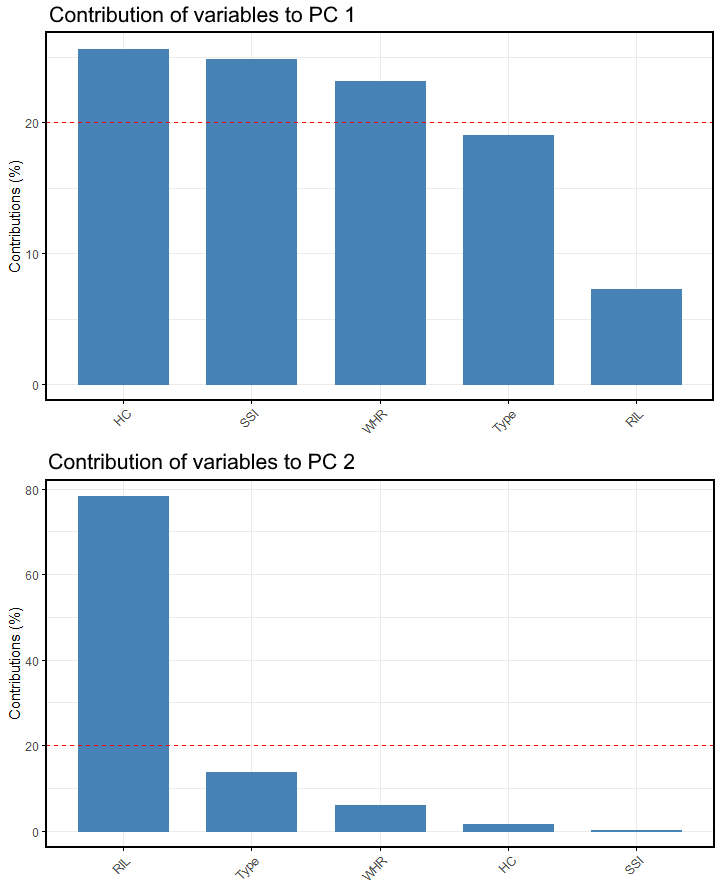

**Supplementary Figure 1**: Contributions of variables to PCs 1 and 2 of PCA run on all quantified ocular traits.

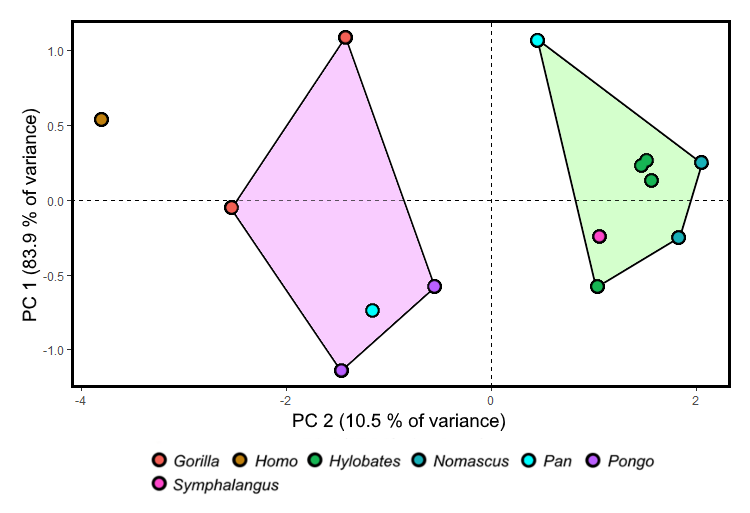

**Supplementary Figure 2**: PCA of ocular traits, excluding RIL. Eigenvalue of PC1 = 3.25, eigenvalue of PC2 = 0.45. Variable contributions are shown in Supplementary Figure 3.

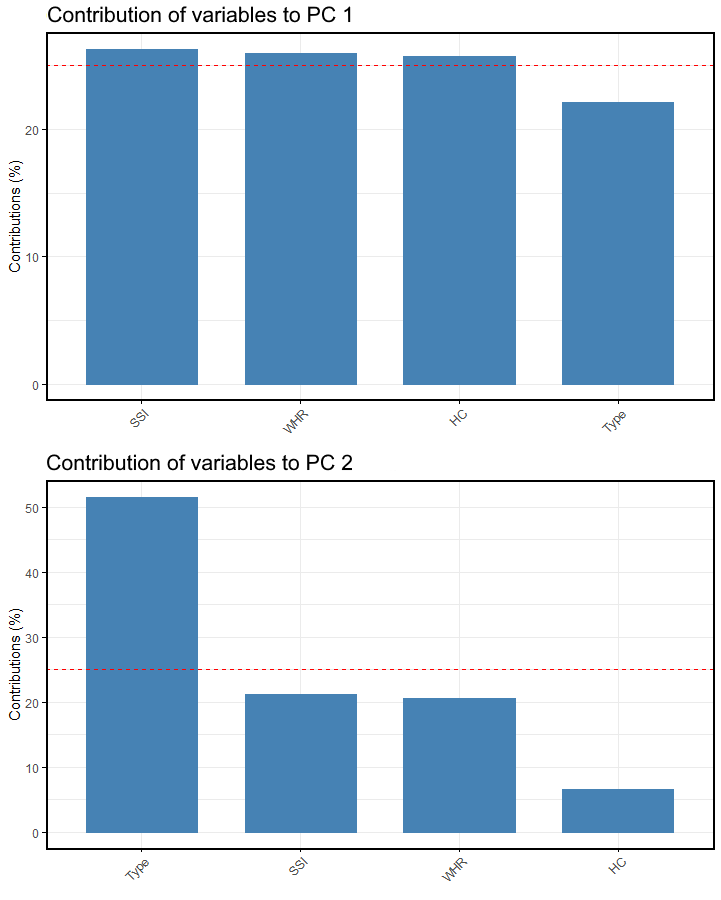

**Supplementary Figure 3:** Contributions of variables to PCs 1 and 2 of PCA omitting RIL.

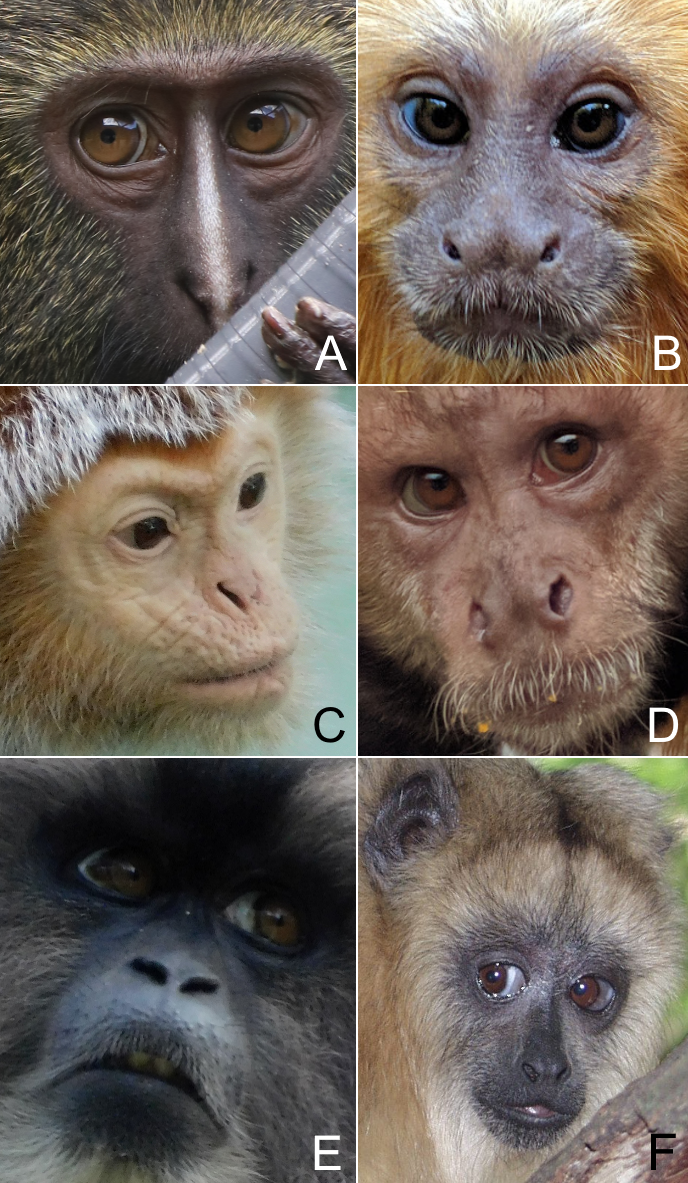

**Supplementary Figure 4:** Pronounced scleral depigmentation in diverse captive New World and Old World monkey species. A: *Cercopithecus hamlyni*, B: *Leontopithecus chrysomelas*, C: *Trachypithecus auratus*, D: *Sapajus xanthosternos*, E: *Macaca silenus*, F: *Alouatta caraya*. Photo credit: A - Miriam Lindenmeier; F - Thomas Geissmann; remaining pictures – Kai R. Caspar.
